# scMultiPreDICT: A single-cell predictive framework with transcriptomic and epigenetic signatures

**DOI:** 10.64898/2026.04.08.717080

**Authors:** Ewura-Esi Manful, Yasin Uzun

## Abstract

Cellular responses to genetic perturbations depend on both transcriptional programs and the epigenetic landscape. While single-cell multiomics technologies enable simultaneous profiling of gene expression and chromatin accessibility, the relative contribution of each regulatory layer to gene expression remains unclear. Existing computational approaches focus on data integration and gene regulatory network inference but do not systematically compare the predictive performance of transcriptional versus epigenetic features on a gene-by-gene basis.

We present scMultiPreDICT, a computational framework for comparative predictive modeling of gene expression using single-cell multiomics data. scMultiPreDICT benchmarks RNA-only, ATAC-only and multimodal feature sets across six machine learning models including regression, tree-based learning and deep learning using multiple biological datasets. We show that RNA-derived features generally provide strong predictive power, whereas chromatin accessibility alone yields a modest performance. Surprisingly, multimodal integration does not uniformly improve prediction accuracy; instead, its benefit is gene-specific and context-dependent. Feature importance analysis reveals that transcriptional features dominate for most genes, whereas chromatin accessibility contributes meaningfully for a subset of genes in specific cellular contexts. Overall, the results demonstrate that regulatory layers contribute differently to gene expression. scMultiPreDICT provides a systematic framework for identifying the relative contributions of transcriptional and epigenetic regulation across genes and cellular contexts, guiding the design of targeted perturbation studies and the prioritization of regulatory layers for therapeutic interventions.

scMultiPreDICT is implemented in R and available at https://github.com/UzunLab/scMultiPreDICT/.

## Introduction

Predicting cellular responses to genetic perturbations and pharmacological interventions remains a fundamental challenge in biology and medicine. Transcriptional programs are widely regarded as the primary driver of a cell’s identity and functional state where key transcription factors are known to control cellular processes (1). However, accumulating evidence challenges this view. Recent studies further demonstrate that the epigenetic landscape can actively shape cellular fate through epigenetic priming in which chromatin accessibility decides the fate of a cell weeks before the activation of corresponding transcriptional programs (2, 3). Additional work has described multiple epigenetic preparation mechanisms including tolerance, reining, priming and transcriptional memory that collectively regulate how cells respond to future perturbations and influence cellular development (4). In some cellular contexts, chromatin states function as a safeguarding mechanism that influences transcriptional programs and restricts cell fate transitions (5, 6).

Chromatin accessibility reflects coordinated regulatory interactions underlying gene expression control (7) and chromatin states within a given cellular context can influence how cells respond to genetic perturbations and reprogramming signals (8). Together, these findings suggest that cellular responses to perturbations such as CRISPR (Clustered Regularly Interspaced Short Palindromic Repeats)-based editing, transcriptional activation, or small molecule inhibition are shaped not only by transcriptional interactions but also by the underlying epigenetic architecture that modulates transcriptional changes.

The emergence of single-cell multiomics technologies, which simultaneously profile gene expression (scRNA-seq) and chromatin accessibility (scATAC-seq) in the same cell, provides an unparalleled opportunity to study these regulatory mechanisms (9). Existing computational approaches predict gene expression from chromatin accessibility (10, 11), infer gene regulatory networks from single-cell multiomics data (12, 13) or integrate transcriptomic and chromatin accessibility for joint analysis and embedding construction (14–16).

Despite these advances, the relative contribution of gene interactions versus cis-regulatory elements in determining gene expression has not been elaborated. This gap limits our ability to identify which regulatory layer is most informative for individual genes and whether multimodal integration provides added value beyond single-modality approaches. Answering these questions is vital for guiding targeted therapeutic design by determining whether to modulate gene regulatory relationships, alter chromatin accessibility or consider both layers jointly.

To address this gap, we introduce scMultiPreDICT (single-cell Multimodal Predictor of Gene Expression via Integrated Chromatin and Transcriptome), a computational framework for systematic gene-by-gene predictive modeling. For each target gene, scMultiPreDICT trains interpretable machine learning models to predict gene expression using three feature sets: 1) expression of other genes derived from scRNA-Seq modality, 2) chromatin accessibility peaks around the transcription start site derived from scATAC-Seq modality, and 3) a combined multimodal feature set integrating features from both modalities. We benchmark a diverse range of models, including baseline linear models, regularized regression methods, tree-based approaches and deep neural networks across multiple biological systems.

Applying scMultiPreDICT to diverse single-cell multiomics datasets spanning developmental and immune contexts, we show that RNA-derived features generally provide strong predictive accuracy, while the contribution of chromatin accessibility is gene-specific and context-dependent. Feature importance analysis reveals key transcriptional regulators and putative regulatory elements that drive gene expression, providing candidate targets for perturbation experiments and identifying potential therapeutic intervention areas. By comparing predictive performance across feature sets, scMultiPreDICT enables the assessment of which regulatory layer is most informative for individual genes and identifies specific features, genes or regulatory elements that contribute to expression variability.

## Methods

### Data acquisition and preprocessing

We used paired scRNA-seq and scATAC-seq data from three distinct biological contexts to evaluate model robustness and generalizability. These included two mouse embryonic stem cell (mESC) datasets at developmental stage ESC (biological replicates 1 and 2) and a human peripheral blood mononuclear cell (PBMC) dataset from 10x Genomics. Quality control was performed independently for each dataset to account for data-specific technical variations using modality-specific thresholds. Low-quality cells were filtered out using RNA and ATAC-specific quality control metrics, with data-specific thresholds chosen based on metric distributions. Detailed filtering criteria for each dataset are provided in the Supplementary Methods (Supplementary Table S1, Figure S1). For the PBMC data, an initial preprocessing and clustering step was performed to isolate a homogeneous cell population. Cell types were annotated using SingleR with the Human Primary Cell Atlas reference (17), and the dataset was subset to include only cells annotated as T cells for downstream analysis. Genes expressed in at least 10% of the filtered, high-quality cells were kept for further downstream analysis.

### Data partitioning

For each dataset, cells were randomly partitioned into a 70% training set, a 20% validation set, and a 10% test set (Supplementary Figure S1). The data split was performed at the cell level prior to any downstream analysis to prevent information leakage between splits.

### Dimensionality reduction and multimodal integrated analysis

For scRNA-seq, gene expression counts were log-normalized and for the scATAC-seq, peak accessibility matrices were normalized using frequency-inverse document frequency (TF-IDF). To obtain a shared representation of scRNA-seq and scATAC-seq for constructing metacells, we evaluated four multimodal integrated analysis strategies. All dimensionality reduction procedures were performed exclusively on the training set with learned representations projected onto the validation and test sets. For unimodal dimensionality reduction, principal component analysis (PCA) was performed on scRNA-seq while latent semantic indexing (LSI) (18) was performed on the scATAC-seq data.

The integrated analysis strategies included PCA+LSI (a linear concatenation of modality-specific embeddings) (16), weighted nearest neighbors (WNN; a neighborhood-based approach that learns and weights the relative contribution of different modalities at the cell level, enabling the integrated analysis of multiple modalities) (19), non-linear latent modeling using single-cell Variational Inference (scVI; a probabilistic framework for analysing scRNA-seq data) (20) for scRNA-seq and PeakVI (a deep generative model designed for single-cell chromatin analysis) (21) for scATAC-seq and joint latent modeling using Multimodal Variational Inference (MultiVI; a probabilistic model that creates a joint representation for analysing multimodal single-cell data) (22).

Throughout this study, PCA+LSI and scVI+PeakVI are treated as paired modality-specific integrated analysis strategies, representing linear and non-linear approaches, respectively. In both cases, embeddings were learned independently for each modality and subsequently z-scored and concatenated to form a combined representation. For all strategies, the resulting low-dimensional representations were used to define k-nearest neighbors (kNN) to construct metacells via kNN smoothing. We refer to these approaches as integrated analysis strategies, as they are used to construct shared neighborhood representations for denoising rather than to directly integrate modalities for prediction. Additional implementation details are provided in the Supplementary Methods section.

### Metacell construction using kNN smoothing

To reduce the technical noise of single-cell data while preserving biological heterogeneity, we implemented kNN smoothing to construct metacells as a denoising strategy. Smoothing was performed separately within the training, validation and test splits to prevent information leakage. For each cell, k nearest neighbors (k = 20, including the cell itself) were identified in the corresponding low-dimensional representation obtained from each integrated analysis using Euclidean distance. Metacell profiles were computed by averaging gene expression counts and chromatin accessibility signals across each cell’s neighbourhood. The resulting metacell-level gene expression counts and chromatin accessibility were normalized using counts per million (CPM) and used as the primary input for all prediction models.

### Feature extraction per target gene

For each target gene, we constructed three distinct feature sets to enable comparison of transcriptional and epigenetic predictors. **RNA-only features** consisted of the expression of the top highly variable genes (HVGs) selected from the training set using a variance-stabilizing transformation. If the target gene was among the selected HVGs, it was excluded from the feature set resulting in 999 predictors. **ATAC-only features** consisted of peaks located within ± 250*kb* of the target gene’s transcription start site (TSS) capturing both proximal promoter regions and distal cis-regulatory elements. To reduce sparsity, only target genes with at least five accessible peaks within this window were retained, giving rise to variable numbers of features across genes. From the initial set of 100 target genes, this filtering excluded genes with fewer than five peaks yielding 96 target genes for mESC replicate-1, 93 genes for mESC replicate-2 and 81 genes for T Cells. The number of peaks per gene ranges from 5 to 125 with a median of 64, 65 and 43 peaks for ESC replicate 1, ESC replicate 2 and T cells respectively substantially lower than RNA-only features. **Multimodal features** were formed by concatenating RNA-only and ATAC-only features for each target gene. Feature set distributions can be found in Supplementary Figure S2.

### Model training and prediction

scMultiPreDICT implements six machine learning models spanning linear models and non-linear approaches. Linear models included ordinary least squares (OLS) (this serves as a baseline linear model without regularization), Ridge regression (L2 regularization), Lasso regression (L1 regularization), and Elastic Net (combined L1 and L2 regularization). Non-linear models included a Random Forest regressor and a deep neural network (DeepNN). The DeepNN was implemented as a multi-layer perceptron neural network using Keras R-interface with three hidden layers using ReLU activation. Model hyperparameters were optimized via a parameter grid search using the validation set. Prior to training, all input features were scaled to [0,1] range using min-max normalization with parameters learned from the training set. Models were trained separately for each target gene. Predictions were performed for two different target gene sets: HVGs and randomly selected non-HVGs. This design enabled us to assess model performance across different gene sets.

### Model evaluation

Model performance was evaluated on the held-out test set using three metrics. The coefficient of determination (*R*^2^) was used to quantify the proportion of variance in the target gene expression explained by the predictors. Root mean squared error (RMSE) was used to measure the average magnitude of prediction errors. In addition, Spearman’s rank correlation coefficient was used to evaluate the relationship between the predicted and actual gene expression. All metrics were computed independently for each target gene, and results were aggregated across approximately 100 target genes to generate performance distributions for comparative analysis across all models and feature sets. For visualization purposes only, quantile normalization was applied to predicted and observed gene expression values when plotting scatter plots for correlation trends. However, all other reported performance metrics were computed on the original values.

### Feature interpretation

To extract biological insights and understand features used by the models for the predictions, we examined the regression coefficients for the linear models and computed feature importance scores (using Gini importance scores) for Random Forest. For models trained on multimodal features, we quantified the relative contribution of each modality by aggregating the absolute coefficient values and feature importance scores separately for RNA-derived and ATAC-derived features. This provided insights into the predictive influence of transcriptional versus chromatin accessibility information for each target gene. Furthermore, due to the unequal numbers of RNA and ATAC features, we defined a **Selection Rate** metric to normalize feature importance by feature availability and mitigate modality dominance due to feature count imbalance. It is calculated as the percentage of top 30 features derived from a specific modality, divided by the average total number of input features for that modality (Equation 1).

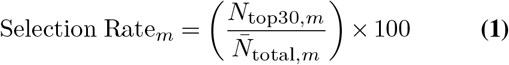

where *m* represents the modality (RNA or ATAC), *N*_top30,*m*_ is the number of features from modality *m* that appear in the top 30 most important features (ranked by importance), 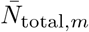 is the average number of available input features for modality *m* across all target genes in the dataset (e.g, the average number of peaks within ± 250 kb for ATAC, or the count of HVGs predictors for RNA).

## Results

### scMultiPreDICT framework and dataset overview

We developed scMultiPreDICT to systematically assess the relative contributions of transcriptional and chromatin accessibility context to gene expression prediction. The framework was applied to three single-cell multiomics datasets: mouse embryonic stem cells (day 7.5, with two biological replicates) and T cells from human PBMCs. Cell quality control was performed independently on each dataset, followed by partitioning into training, validation and test sets. To reduce the sparsity nature of single-cell data, prediction was performed with metacells, which were constructed with dimensionality reduction and kNN smoothing.

For each target gene, three feature sets were evaluated: RNA-only derived from top 1000 HVGs, ATAC-only consisting of peaks within ±250*kb* of the target gene’s TSS and multimodal feature set combining RNA and ATAC features. Predictions were performed for two categories of target genes; HVGs and Non-HVGs. scMultiPreDICT trains six machine learning models for each feature set including OLS, Lasso, Ridge, Elastic Net, Random Forest and DeepNN and evaluates model performance on the held-out test set using Spearman’s rank correlation, *R*^2^ and RMSE on the test set (Figure 1).

**Fig. 1.**
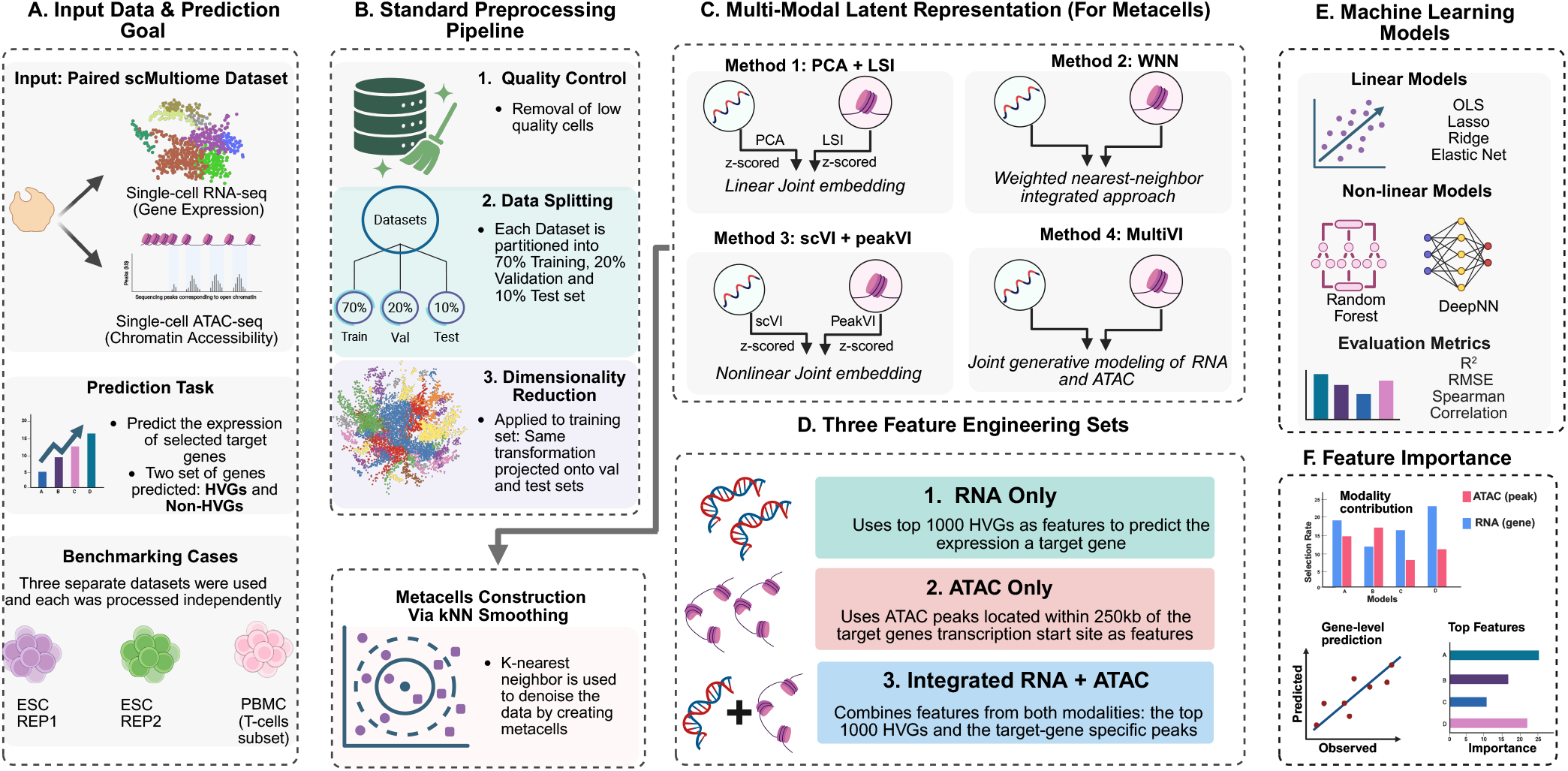
Overview of scMultiPreDICT framework. Schematic workflow for gene expression prediction using single-cell multiome data, including quality control, data splitting, multimodal integrated analysis for metacell construction, feature engineering for RNA-only, ATAC-only and multimodal prediction, model training and prediction and feature interpretation.

### Interaction-based gene expression prediction

We first assessed how well transcriptional data (RNA-derived features) predict individual target gene expressions. Across all datasets and target genes, models with only RNA-based features achieved a high predictive performance (Figure 2A, D). Across the samples, we observed a notable difference between two replicates for ESC. The prediction accuracy was notably higher for the first replicate (Rep1, correlation ranging from 0.66 to 0.78) compared to the second replicate (Rep2, correlation ranging from 0.49 to 0.63). The accuracy difference across two replicates from the same biological sample can possibly be attributed to the overall data quality between the replicates. Although the second replicate contains a larger number of cells (Rep2: n = 8,333) than the first replicate (Rep1: n = 6,484), quality control analysis revealed that the first replicate exhibited better signal quality for both data modalities, including higher median UMI (unique molecular identifier) counts, greater number of detected genes and peaks per cell (Supplementary Figure S2). These quality differences likely influenced the signal-to-noise ratio available for model training, thereby affecting the predictive performance on the unseen (test) cells. Interestingly, the accuracy for the T cell population from the PBMC data (median correlation ranging between 0.60 to 0.75) was in between the two mESC replicates, indicating the prediction accuracy does not necessarily depend on the cell type.

**Fig. 2.**
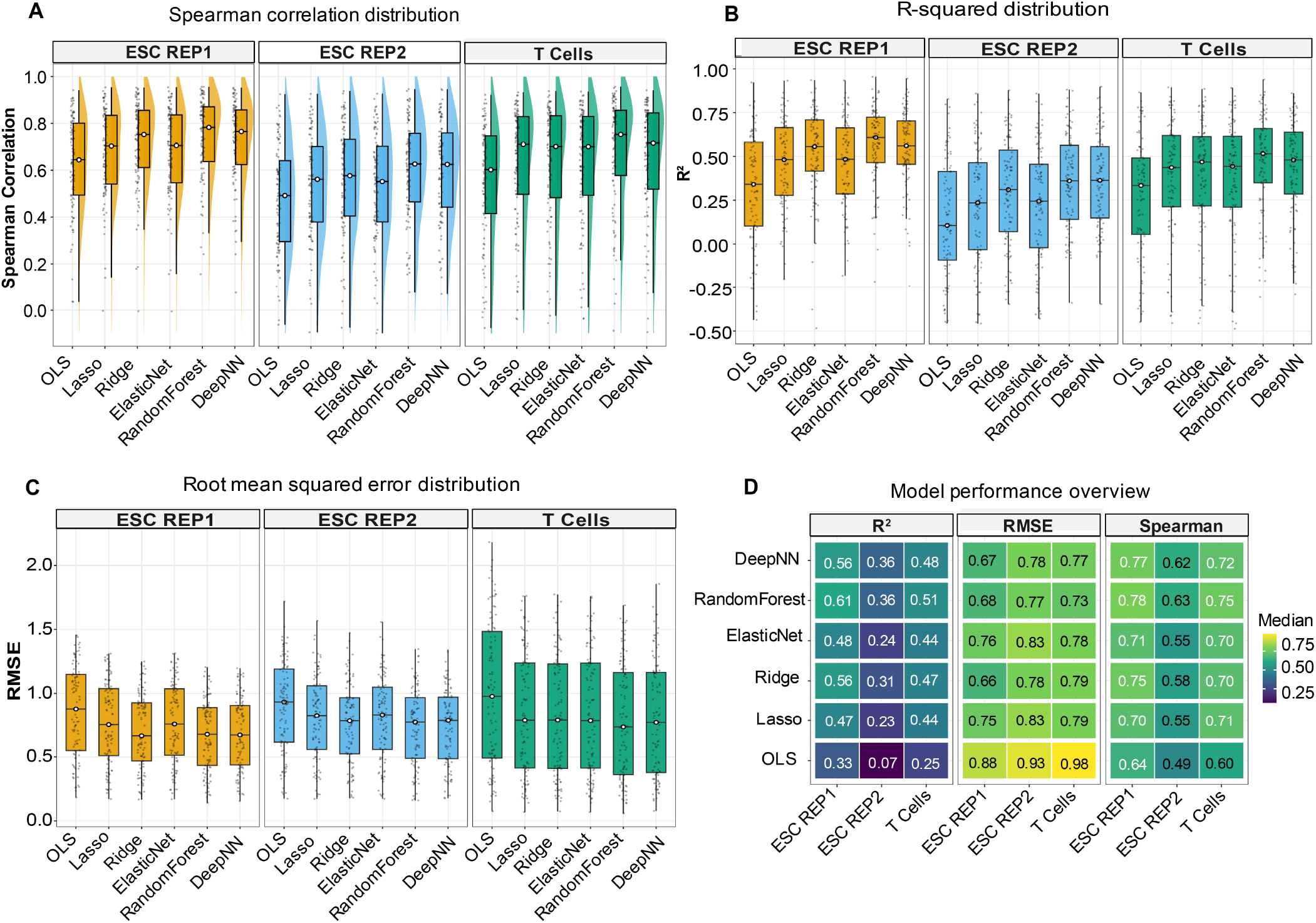
RNA-only gene expression prediction performance. Distribution of prediction performance across models and datasets using RNA-only as features. (A) Spearman rank correlation between predicted and observed expression values. (B) Coefficient of determination (*R*^2^). (C) Root mean squared error (RMSE). D) Heatmap summarizing median performance metrics across datasets and models. Color intensity indicates performance (darker = lower performance, lighter values = higher performance). Results are shown for mESC replicates and T cells from the PBMC dataset in panels A, B and C, with each point representing an individual target gene.

Across all datasets, Random Forest, regularized linear models (Lasso, Ridge, Elastic Net) and DeepNN consistently out-performed ordinary least squares (OLS) (Figure 2A-C). Random Forest achieved the highest overall performance while Deep Neural Network (DeepNN) generally exceeded the regularized models, possibly due to their capabilities in capturing non-linear regulatory associations among the genes. Prediction performance was generally observed for highly variable genes (HVGs) compared to non-HVGs (Supplementary Figure S3) as the low variation in the response variable reduces learnable signal (signal-to-noise) and instability in the performance metric (loss) during training. Overall, these results demonstrate that the expression of other genes provides significant power for target gene expression prediction, highlighting the interconnected nature of gene regulatory networks underlying cellular function.

### Chromatin accessibility signature-based gene expression prediction

To assess the contribution of cis-regulatory elements in predicting gene expression, we evaluated ATAC-only models using peaks within the (± 250*kb*) flanking region of each target gene’s TSS. Considering all the input datasets and methods, predictions based on chromatin accessibility data achieved moderate performance (median correlation ranging from 0.38 to 0.60, Figure 3A, D). Consistent with the results using features from expression data, mESC replicate-1 exhibited the highest predictive performance (median 0.56-0.60) while mESC replicate-2 and T cells showed considerably lower correlations (median 0.38-0.42). As in the case of expression-only features, prediction accuracy was higher for HVGs compared to non-HVGs (Supplementary Figure S4). Notably, performance differences across models were minimal (Figure 3A-C) when compared to expression-based models. This may be attributed to the lower number of features (peaks per target gene ranging from 5 to 125 with a median of 61, 62 and 42 peaks for mESC Rep1, mESC Rep2 and T cells respectively) compared to the RNA-only features (1000 HVGs), which reduces overfitting risk and thereby limits the advantages of regularization or non-linear modeling. Overall, chromatin accessibility (ATAC)-based models consistently underperformed expression (RNA)-based models in terms of accuracy.

**Fig. 3.**
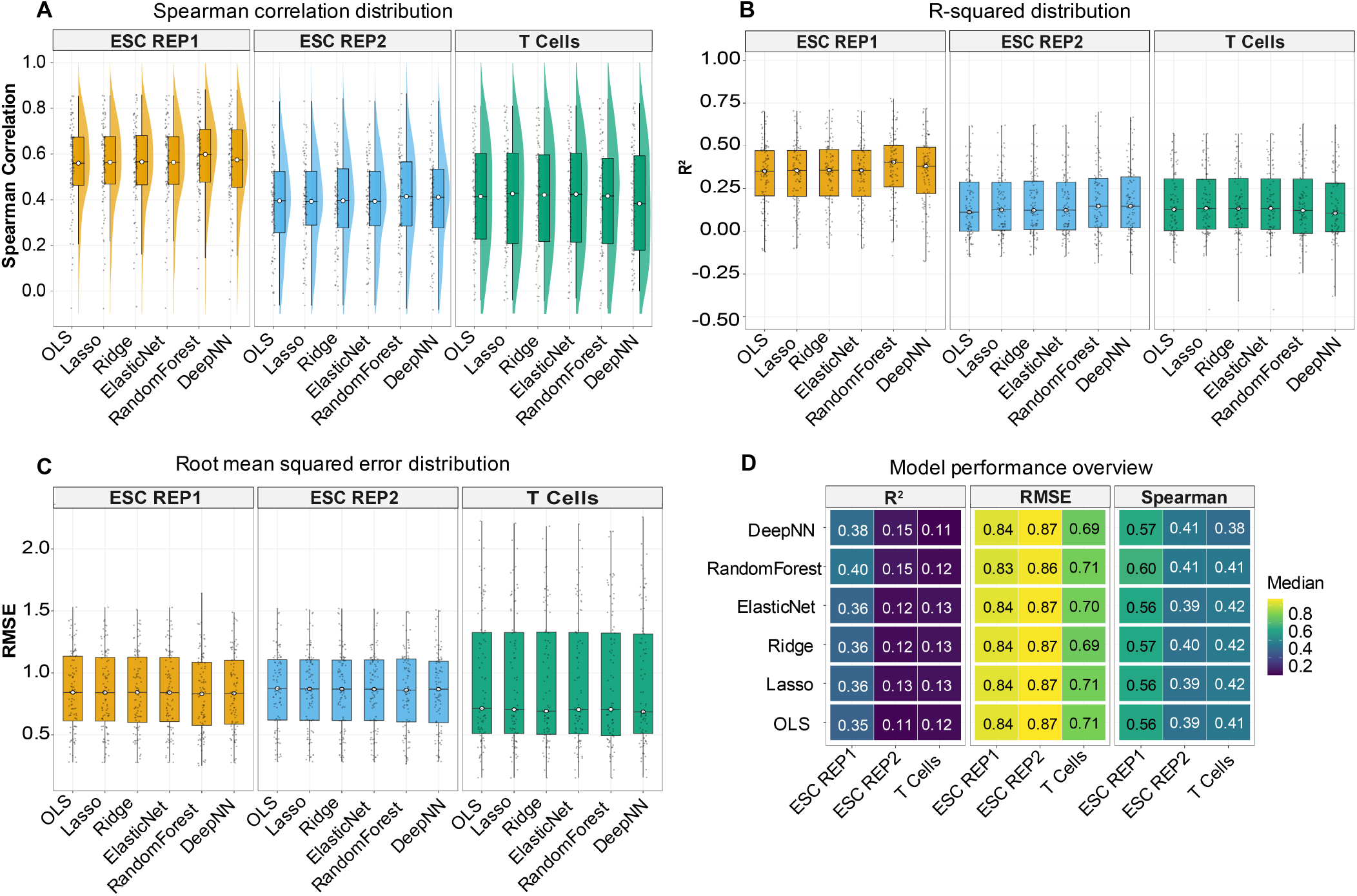
ATAC-only gene expression prediction performance. Distribution of prediction performance across models and datasets using peaks within ± 250*kb* of each target gene’s TSS. (A) Spearman rank correlation between predicted and observed expression values. (B) Coefficient of determination (*R*^2^). (C) Root mean squared error (RMSE). D) Heatmap summarizing median performance metrics across datasets and models. Color intensity indicates performance (darker = lower values, lighter values = higher values). Results are shown for mouse ESC biological replicates and PBMC T cells, with each point representing an individual target gene.

### Multimodal predictive performance

Given that RNA-only features capture gene regulatory networks, and ATAC-only features capture its contribution to gene expression, we hypothesized that combining both modalities would improve prediction accuracy. We first evaluated multimodal predictions using a PCA+LSI integrated analysis in which linear dimensionality reduction was applied separately to scRNA-seq and scATAC-seq and the resulting embeddings were concatenated prior to metacell construction.

With this initial approach, across the tested datasets, multimodal models achieved predictive performance values that are comparable to those observed in expression features-only prediction (Figure 4A). Contrary to our initial expectation, the multimodal prediction with the inclusion of chromatin accessibility features did not consistently outperform RNA-only models across target genes or datasets. Performance improvement was gene-dependent (Supplementary Figure S7), with a subtle overall improvement. This result indicates that the predictive contribution of chromatin accessibility is not universal and regardless of the dataset and the method utilized, the synergistic effect was not observable when the set of genes is considered as a whole.

**Fig. 4.**
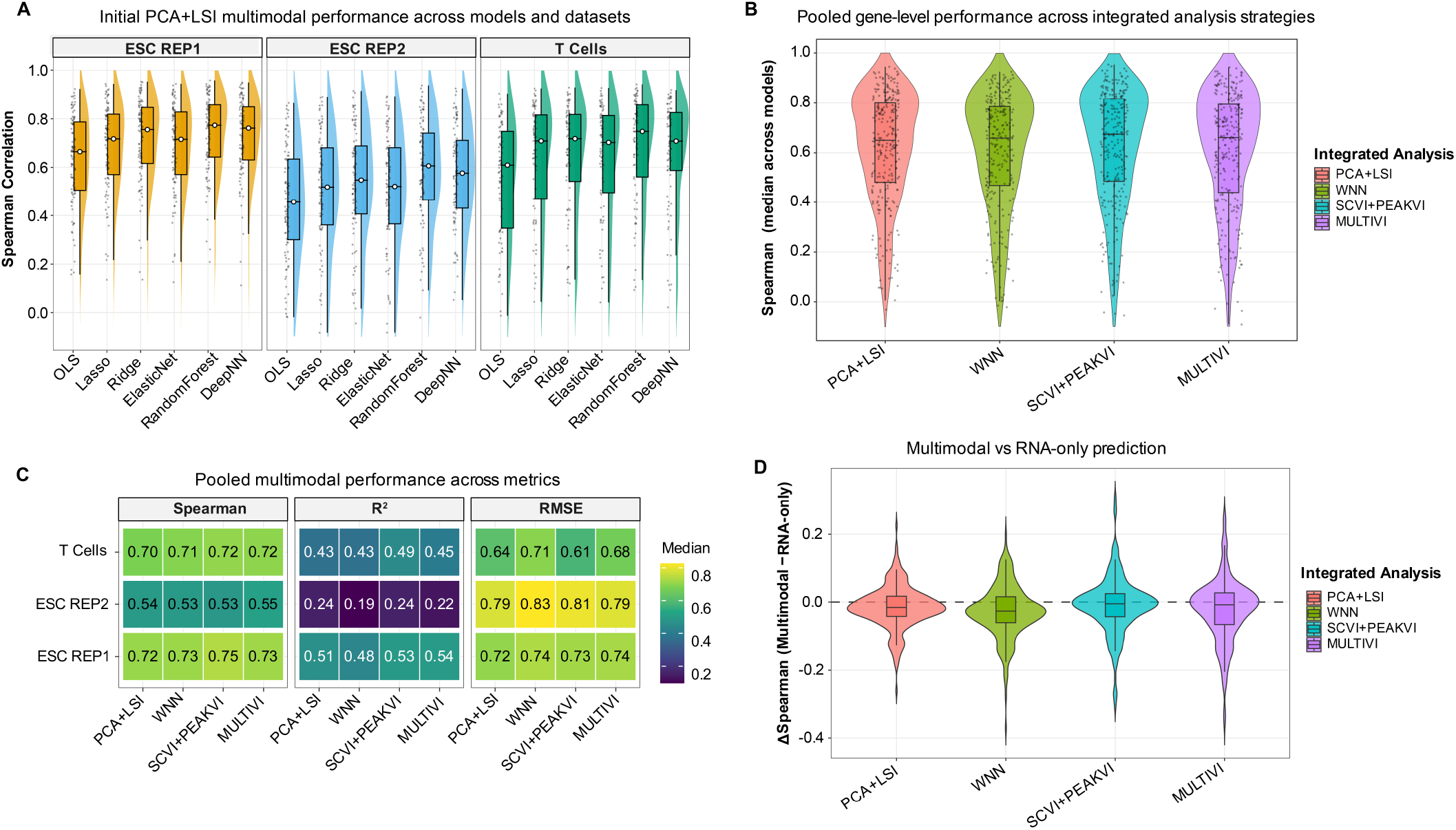
Multimodal gene expression prediction across integrated analysis strategies. Performance of multimodal (RNA+ATAC) prediction under different integrated analysis approaches. (A) Spearman correlation distributions across models and datasets using the initial PCA+LSI integrated analysis. (B) Pooled gene-level Spearman correlations across all models and datasets comparing four integrated analysis strategies (PCA+LSI, WNN, scVI+PeakVI, and MultiVI). (C) Heatmap summarizing median pooled performance across integrated analysis strategies and datasets for Spearman correlation, *R*^2^, and Root mean squared error (RMSE). (D) Difference in predictive performance between multimodal and RNA-only models (ΔSpearman), highlighting the relative increase or decrease from multimodal integration. Each point represents an individual target gene.

We therefore explored whether alternative integrated analysis strategies that can better extract shared embedding space between the two modalities could enhance the prediction accuracy. We employed four embedding strategies for this purpose: PCA+LSI, WNN, scVI+PeakVI and MultiVI. To enable a direct comparison at the gene-level, prediction metrics were pooled across all models and datasets for each integrated analysis. Comparing four strategies from the pooled metrics, we observed no improvement in the aggregated gene-level predictive performance over RNA-only prediction as reflected by the similar correlation distributions and closely related median performance metrics (Figure 4B-C). Despite differences in underlying modeling assumptions, none of the integrated analysis strategies consistently outperformed the models with expression-only features (Figure 4D)

We further performed gene-level analyses to determine whether the observed underperformance of multimodal models relative to expression-only feature models was uniform across target genes. We found that while a subset of genes exhibited improved predictive performance when chromatin accessibility features were added, many showed little or no change and some showed reduced performance relative to RNA-only models (Supplementary Figure S7). Classification of target genes based on performance improvement revealed that only a limited subset of genes consistently benefited from multimodal integration across strategies (Supplementary Figure S7; Supplementary Table S2). Together, these findings demonstrate that multimodal predictive performances are gene-specific and are not primarily driven by the choice of the integrated analysis strategy.

### Feature importance analysis reveals key regulators and cis-regulatory elements

To interpret the key drivers of the multimodal predictions, we measured the feature importance by quantifying the composition of the top (30) predictive features in terms of category (ATAC:peaks or RNA:genes). Because the two feature sets differ substantially in size, we used an adjusted metric to mitigate the bias caused by the high number of expression-based features relative to the number of proximal peaks.

The composition of the top features was dependent on the cell type (Supplementary Figure S8). For both mESC datasets, RNA-derived features dominated the top predictors for most target genes whereas in T cells, ATAC-derived features contributed at levels comparable to RNA-derived features, indicating increased contribution from chromatin accessibility in this cellular context (Supplementary Figure S8).

Using results from the best performing model, Random Forest, we examined three representative target genes in detail, one from each dataset to explore the diversity of regulatory mechanisms specific to each cell type. The genes *Etv6* and *Tbx3* both from mESC replicates showed predictions primarily driven by RNA-derived features. The results revealed that *Etv6* top predictors include *Pbx1, Cbx5* and *Lin28b* while *Tbx3* depended on *Prdm6, Dlk1* and *Polg*. In contrast, *RUNX3*, selected from the T cells dataset, showed contributions from both RNA-derived features and ATAC-derived features indicating an involvement of cis-regulatory elements. The top predictors include *TSHZ2*, a local ATAC peak, chr1:24910685-24911687 followed by RNA features including *LEF1* and *FHIT* (Supplementary Figure S8). These examples illustrate the diversity of regulatory feature combinations identified by scMultiPreDICT.

Feature importance rankings from Random Forest identified specific transcriptional and proximal accessible chromatin regions associated with each target gene (Supplementary Figure S8), providing candidate targets for experimental validation. These gene-specific patterns further support the idea that chromatin accessibility selectively contributes to gene expression predictions and that multimodal integration enables the elucidation of regulatory mechanisms across different cell types.

## Discussion

In this study, we present scMultiPreDICT, a gene expression prediction framework for quantifying transcriptional and epigenetic contributions to gene expression using single-cell multiome data. By systematically benchmarking RNA-only, ATAC-only and multimodal prediction across multiple machine learning models and biological datasets, scMultiPreDICT enables interpretable assessment of regulatory influences beyond descriptive multimodal analysis.

We observed that transcriptional-based information provides strong predictive accuracy across datasets, particularly for highly variable gene expression. This outcome likely reflects the coordinated nature of gene expression, where the expression of individual genes is influenced by broader gene-gene interactions within a cell. In contrast, chromatin accessibility alone yields moderate predictive performance compared to transcriptional features.

The relatively weaker predictive accuracy can be attributed to multiple reasons. First, chromatin accessibility is required, but not sufficient for gene expression. In other words, accessibility is permissive for transcription, but not deterministic by itself alone. In addition to accessibility, multiple other epigenetic modulators can affect gene expression, including DNA methylation and histone modification, which cannot be inferred using ATAC-Seq profiling. Second, accessibility precedes transcription, with a time lag between the two events. Hence, for those genes for which the regulatory elements have recently changed their accessibility status, this update cannot be reflected for the transcription at the time of sample collection. This time lag in the transcription process may have caused a temporal discrepancy between the two modalities.

Motivated by the complementary functions of transcriptional and epigenetic regulation, we evaluated the accuracy of multimodal prediction by combining RNA-derived and ATAC-derived features. We found that while multimodal models achieved strong predictive performance, they did not consistently outperform RNA-only models. Interestingly, this pattern was consistent between different integrated analysis strategies, suggesting that differences in integrated analysis strategies have limited impact on the general accuracy of the prediction.

Among the machine learning models evaluated, Random Forest consistently achieved the highest predictive performance across all datasets, followed by neural network (DeepNN), regularized regression methods (Lasso, Ridge, and Elastic Net) and OLS. The strong performance of Random Forest likely reflects its ability to capture non-linear relationships and feature interactions highlighting the inherent non-linear nature of regulatory processes underlying cellular mechanisms. While neural networks demonstrated comparable performance by modeling complex non-linear patterns, they generally benefit from larger sample sizes in order to generalize well and require extensive hyperparameter tuning. In contrast, regularized linear models achieved moderate performance by mitigating multicollinearity and overfitting but were constrained by their linear assumptions, while OLS underperformed possibly due to its lack of regularization, inability to capture non-linear interactions and sensitivity to high dimensional predictor space.

Gene-level analysis further revealed that the benefit of the multimodal approach is gene-specific. While a subset of genes exhibited improved accuracy with the inclusion of chromatin accessibility features, many showed little or no improvement with some having reduced performance relative to the RNA-only models. Feature importance analysis indicated that RNA-derived features dominate prediction for most target genes, whereas ATAC-derived features contribute meaningfully for a subset of genes, particularly in specific cellular contexts.

By examining the top predictive features for representative target genes, scMultiPreDICT provided insights into the biological basis of the prediction patterns observed. *Etv6* and *Tbx3* from the embryonic stem cell were primarily associated with transcriptional regulators, which is consistent with prior work highlighting their central roles in transcriptional control of pluripotency and differentiation. *Tbx3* is a well-characterized transcriptional regulator required for maintaining self-renewal capacity and regulating differentiation programs in mouse embryonic stem cells (23) while *Etv6* has been shown to play an important role in hematopoietic lineage commitment (24). In contrast, *RUNX3* in T cells exhibited a mixed regulatory signature with a local peak as its second top predictor alongside the transcription factors *TSHZ2* and *LEF1*, illustrating an example where both regulatory layers may be important for a cell’s state. This observation is consistent with *RUNX3*’s established role in T cell development and lineage specification (25). *RUNX3* is known to function through cooperative interactions with other transcription factors to regulate T cell homeostasis and function (26, 27) supporting our observed top predictors. Moreover, long-range enhancer-specific interactions at the *Runx3* locus during T cell development have been reported in mice (28) supporting a role for chromatin organization in controlling *RUNX3* expression. The functional role of these top predictors warrants further experimental validation.

Together, these results indicate that while gene expression features are generally the strongest predictors of target gene expression, specific genes and distinct cell types may exhibit increased contribution from chromatin accessibility. The identification of top predictive features has practical implications for perturbation studies where the transcriptional responses of genetic manipulation may be dependent on the epigenetic landscape. This would influence the prioritization of targets for which the modulation of chromatin accessibility is more likely to influence gene expression as opposed to genes whose expression may depend primarily on the transcriptome.

## Acknowledgements

The authors acknowledge computational resources provided by the Pennsylvania State University College of Medicine.

## Author contributions

E.E.M did the analytical work. Y.U supervised the study.

E.E.M and Y.U wrote the manuscript together.

## Data availability

Publicly available datasets were used in the study. The PBMC data is available from the 10x Genomics dataset repository and can be found at https://www.10xgenomics.com/datasets/10-k-human-pbm-cs-multiome-v-1-0-chromium-x-1-standard-2-0-0 (29). The Mouse Embryonic Stem Cell (mESC) is available from the Gene Expression Omnibus (GEO) under accession number GSE205117 (30). Specifically, we analyzed the E7.5 developmental stage (Replicates 1 and 2).

