## Supplementary Materials (Supplementary Document and Table S2) for "scMultiPreDICT: A single-cell predictive framework with transcriptomic and epigenetic signatures": Supplementary.pdf

#### Methods (Expanded Version)

##### Quality Control and Data Processing

Cells were filtered separately for each dataset using RNA- and ATAC-specific quality control metrics. Thresholds were selected based on the distributions of quality metrics within each dataset to remove low-quality cells. The same filtering principles were applied across datasets, but numerical cutoffs varied to account for dataset-specific sequencing depth and technical variability (Supplementary Table S1, Supplementary Figure S1). All downstream analyses were performed on cells passing these dataset-specific quality control criteria. In this context, although ESC replicate-2 contains more cells overall, ESC replicate-1 exhibits higher quality cells across all quality control metrics (Supplementary Figure S2).

**Table S1. Data-specific quality control thresholds and cell counts**

| Quality Control Metric | ESC REP1 | ESC REP1 | PBMC |
|---|---|---|---|
| Minimum RNA Features | 1000 | 1000 | 500 |
| Maximum RNA Features | 7000 | 6000 | 3200 |
| Maximum mitochondrial percentage (%) | 25 | 35 | 15 |
| Minimum ATAC fragment count | 4000 | 1000 | 4000 |
| Minimum RNA count | 100000 | 100000 | 500 |
| Maximum RNA count | 60000 | 40000 | 9000 |
| Maximum nucleosome signal | 1.8 | 1.7 | 1.2 |
| Minimum TSS enrichment | 4 | 3 | 4 |
| Minimum gene detection (%) per cell | 10 | 10 | 10 |
| **Cells before filtering** | 8936 | 13181 | 10970 |
| **Cells after filtering** | 6484 | 8333 | 8763 |

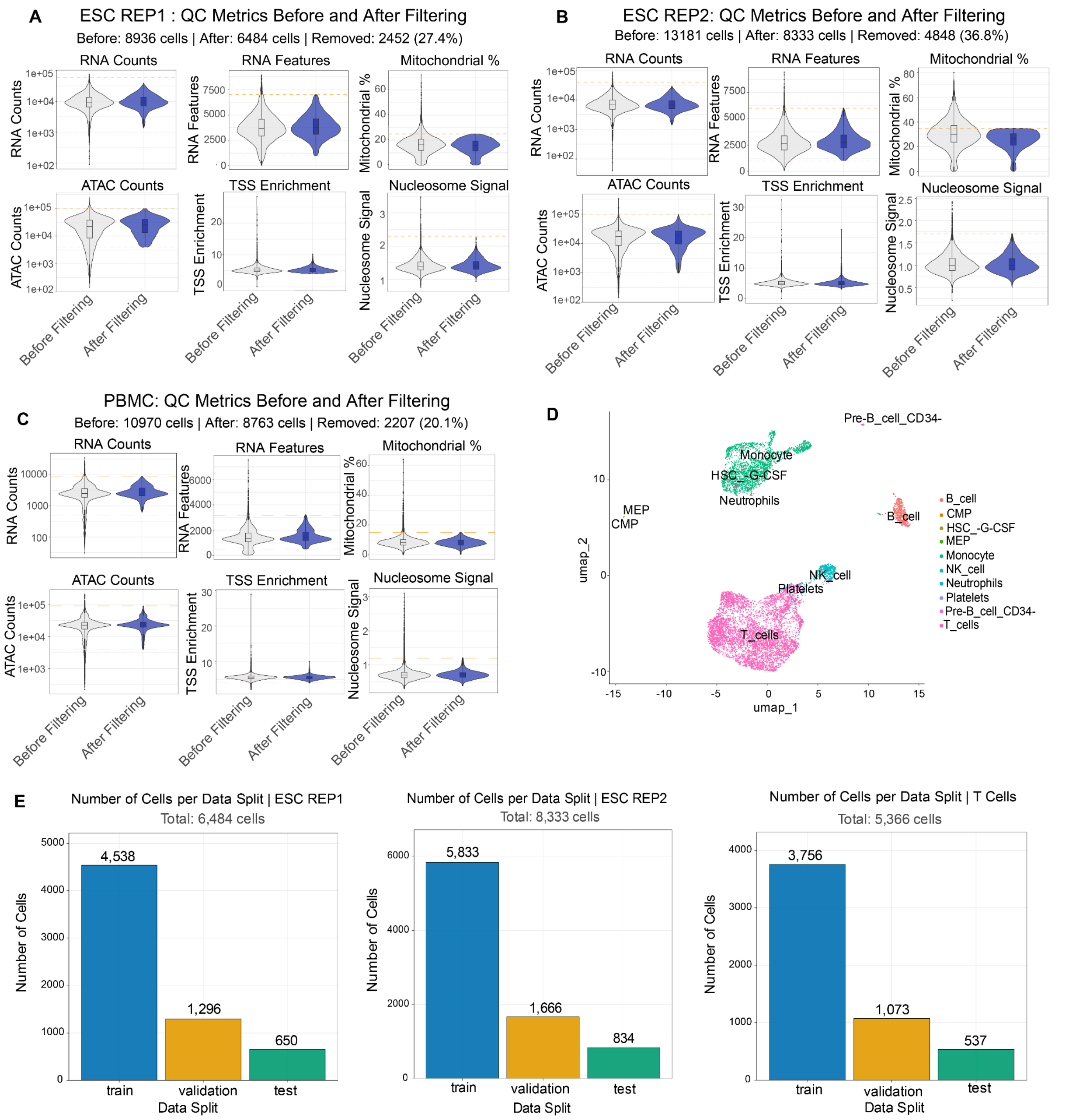


**Supplementary Figure S1: Quality Control, filtering and dataset split composition**.

(A-C): Distributions of RNA and ATAC quality control metrics before and after filtering for ESC replicates 1 and 2, and PBMC datasets, respectively. Metrics include RNA counts, detected RNA features, mitochondrial read percentage, ATAC fragment counts, transcription start site (TSS) enrichment and nucleosome signal. Dataset-specific thresholds were applied to account for differences in sequencing quality.

(D) UMAP visualization of PBMC dataset after filtering, colored by annotated cell types showing clear separation of clusters.

(E) Number of cells assigned to training, validation, and test splits for each dataset following quality control.

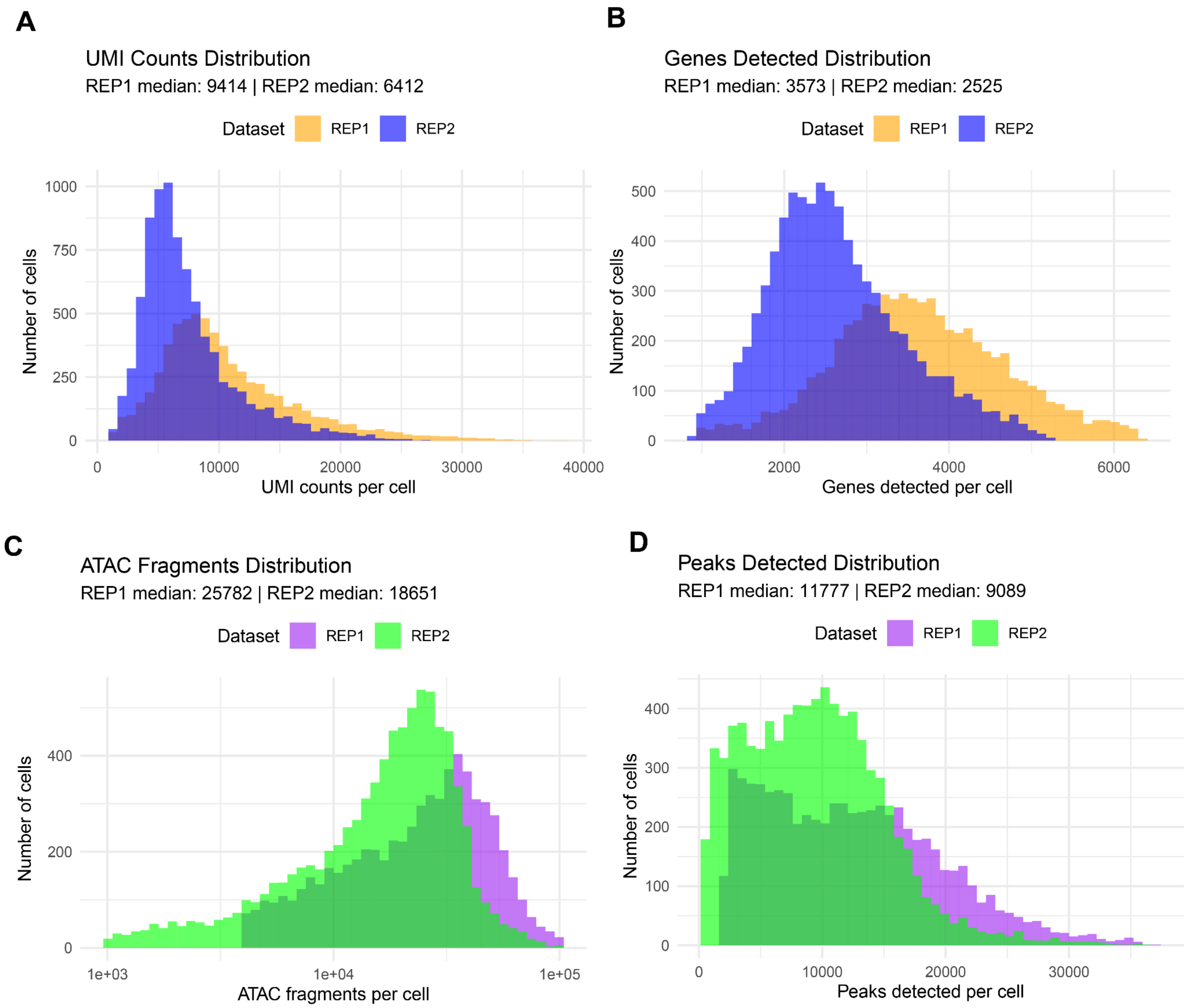


**Supplementary Figure S2: Exploratory comparison of RNA and ATAC signal distributions between ESC biological replicates after cell quality control filtering.** (A) Distribution of RNA UMI Counts per cell for scRNA-Seq data. (B) Distribution of genes detected per cell for scRNA-Seq data. (C) Distribution of fragments per cell for scATAC-Seq data. (D) Distribution of peaks detected per cell for scATAC-Seq data. Median values are noted for each replicate in the title.

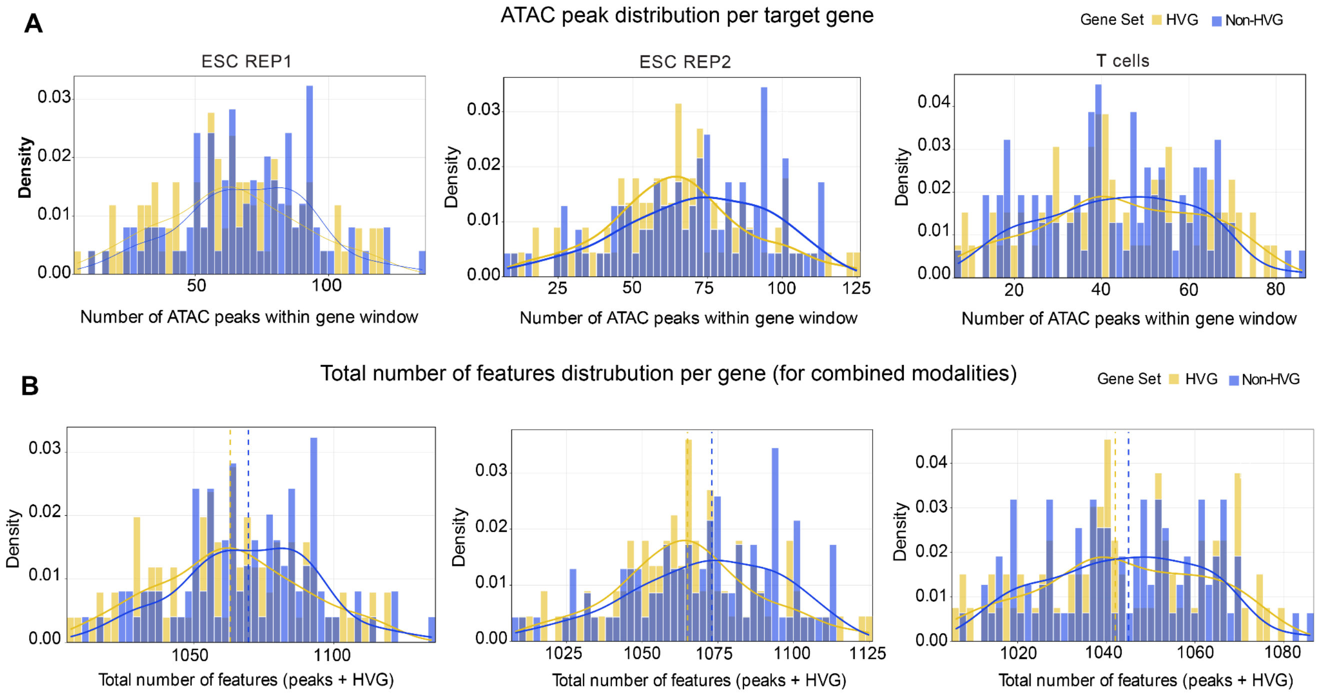


**Supplementary Figure S3: Distribution of ATAC and multimodal feature counts per target gene.** (A) Distribution of the number of chromatin accessibility peaks in the scATAC-Seq data, within ±250kb of each target gene's TSS across ESC replicates 1 and 2, and T cells (PBMC) showing separately for HVG and non-HVGs. (B) Distribution of the total number of features per target gene in the multimodal feature set, combining scRNA-Seq-derived features (1000 HVGs) with gene-specific chromatin accessibility peaks. The dashed vertical lines indicate median feature counts. Feature dimensionality varies across genes and datasets.

**Dimensionality reduction and multimodal integrated analysis**

For scRNA-seq, raw counts were log-normalized and highly variable genes were identified. For scATAC-seq, peak accessibility matrices were normalized using term frequency–inverse document frequency (TF-IDF) as implemented in the Seurat (Hao *et al.*, 2024) and Signac frameworks (Stuart *et al.*, 2022). For unimodal dimensionality reduction, log-normalized scRNA-seq data were reduced using principal component analysis (PCA), while TF-IDF-normalized scATAC-seq data were reduced using latent semantic indexing (LSI).

Four integrated analysis strategies were evaluated to find a shared embedding space between scRNA-seq and scATAC-seq data for the purpose of metacell construction via k-nearest neighbor (kNN) smoothing. All dimensionality reduction models were trained exclusively on the training set, and learned representations were projected onto validation and test sets to prevent information leakage.

The first strategy, PCA + LSI, applied principal component analysis (PCA; 30 components) to scRNA-seq data using the top 3,000 highly variable genes (HVG) and latent semantic indexing (LSI; 30 dimensions) to scATAC-seq data using all peaks. The first LSI component, which primarily reflects sequencing depth, was excluded. Reduced dimensions from both modalities were z-scored and concatenated.

The second strategy applied weighted nearest neighbor (WNN) analysis to modality-specific PCA and LSI embeddings using the Seurat framework, integrating modalities at the level of cell–cell neighborhood graphs with adaptive modality weighting.

The third strategy employed single-cell Variational Inference (scVI) (Lopez *et al.*, 2018) and PeakVI ((a deep generative model designed for single-cell chromatin analysis) (Ashuach *et al.*, 2022) to learn separate 30-dimensional nonlinear latent representations for scRNA-seq and scATAC-seq data from the training set. Latent embeddings were projected onto validation and test sets, z-scored, and concatenated.

The fourth strategy used Multimodal Variational Inference (MultiVI) (Ashuach *et al.*, 2023) to learn a single joint 30-dimensional latent representation directly from the single-cell multiomic training data, which was then used to infer representations for all data splits.

For all integrated analysis strategies, the resulting low-dimensional representations were used exclusively to define k-nearest neighbor graphs for metacell construction via kNN smoothing. As in the main manuscript, these approaches are referred to as integrated analysis strategies, as they are used to construct neighborhood representations for denoising rather than as direct input features for prediction.

### Results (Expanded Version)

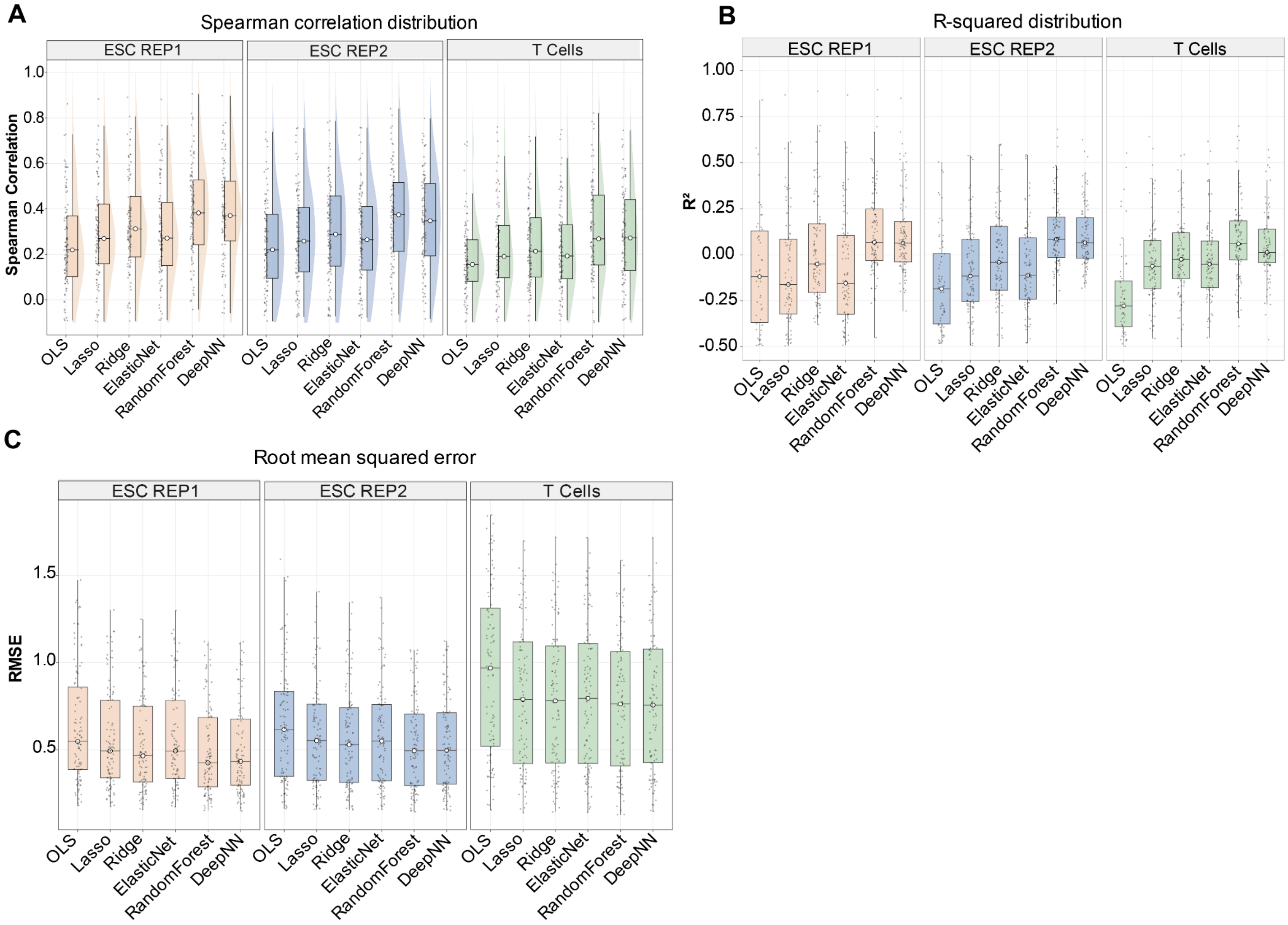


**Supplementary Figure S4: RNA-only gene expression prediction performance for Non-HVG Gene Set.** Distribution of prediction performance across models and datasets using RNA-only as features. (A) Spearman rank correlation between predicted and observed expression values. (B) Coefficient of determination ($R^2$). (C) Root mean squared error (RMSE).

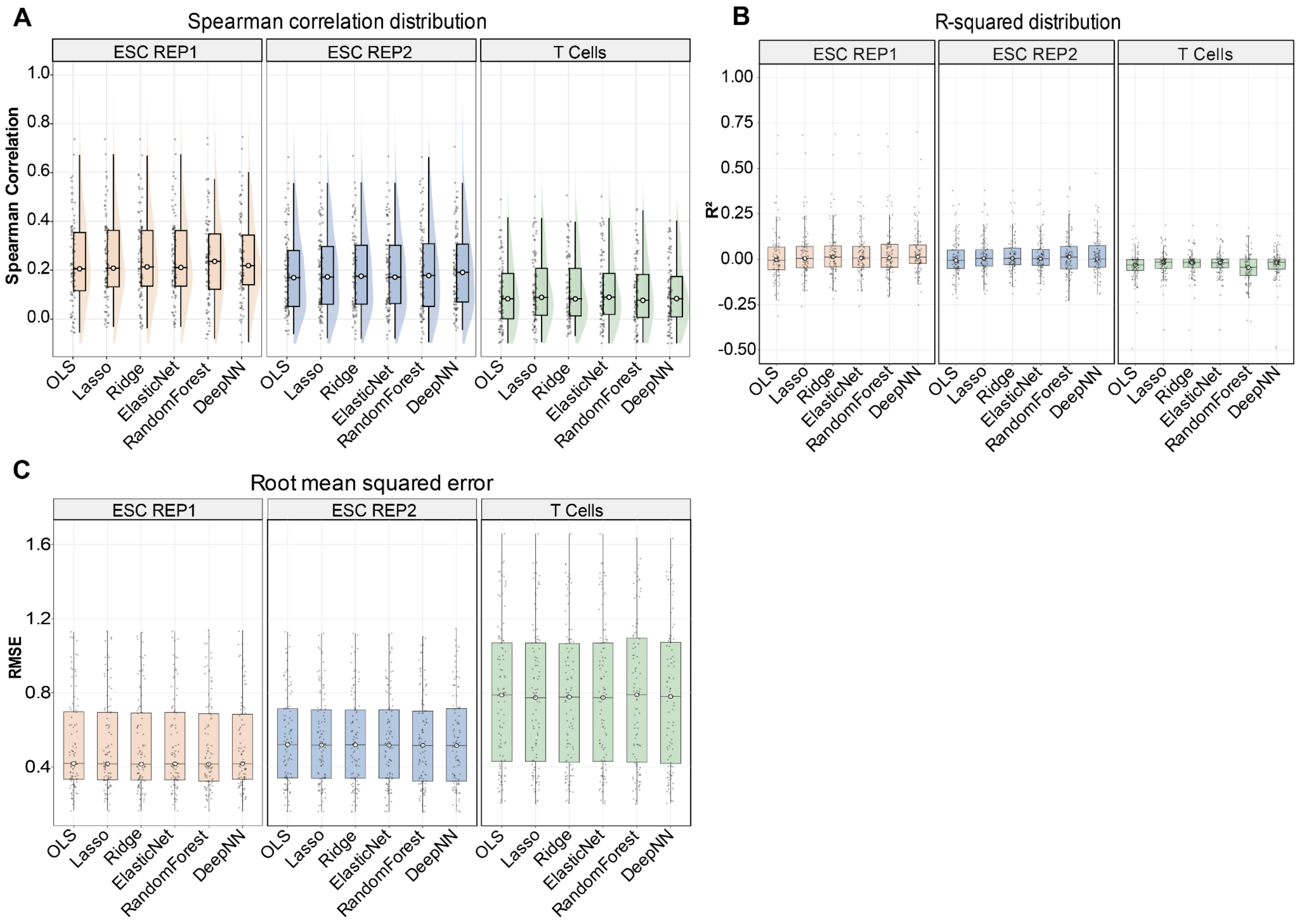


**Supplementary Figure S5: ATAC-only gene expression prediction performance for Non-HVG Gene set.** Distribution of prediction performance across models and datasets using peaks within ±250kb of each target gene's TSS. (A) Spearman rank correlation between predicted and observed expression values. (B) Coefficient of determination ($R^2$). (C) Root mean squared error (RMSE).

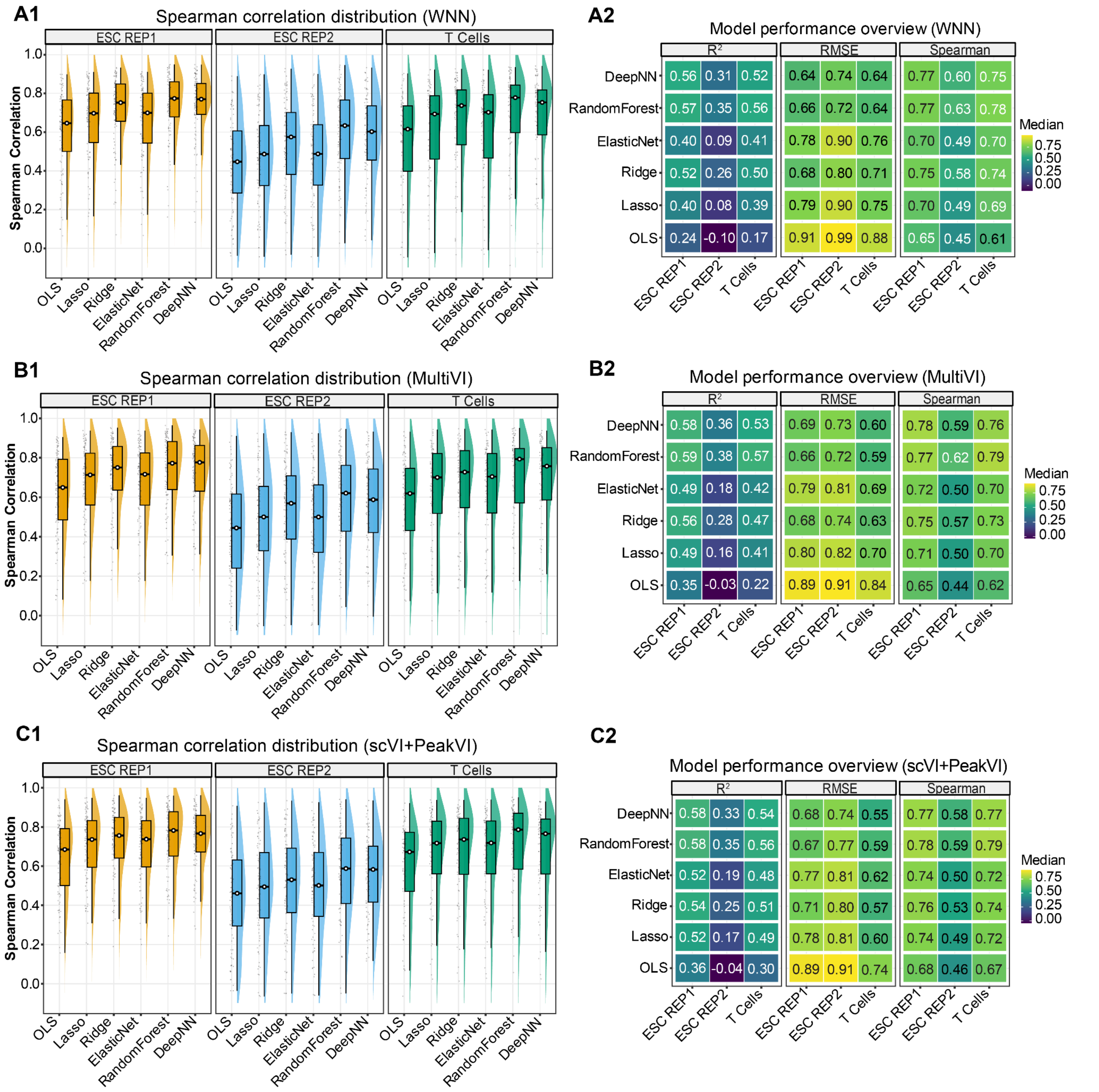


**Supplementary Figure S6: Multimodal prediction performance across alternative integrated analysis strategies for HVG Gene set.**

(A1-C1) Distributions of Spearman Correlations for multimodal prediction using weighted nearest neighbors (WNN), MultiVI and scVI+PeakVI embeddings shown across ESC REP1, ESC REP2 and PBMC T cells for all evaluated models.

(A2-C2) Corresponding model performance overview for each integrated analysis strategy reporting $R^2$, RMSE and Spearman Correlation across datasets and models.

**Multimodal predictive performance**

To assess gene-specific effects of multimodal integration, we computed the difference in Spearman correlation between multimodal and RNA-only prediction (ΔSpearman) for each target gene. Genes were classified as improved (ΔSpearman > 0.01), worse (ΔSpearman < −0.01), or unchanged (|ΔSpearman| ≤ 0.01). Gene-level classifications for all target genes are reported in Supplementary Table S2 and aggregated proportions are summarized for each integrated analysis strategy in Supplementary Figure S7.

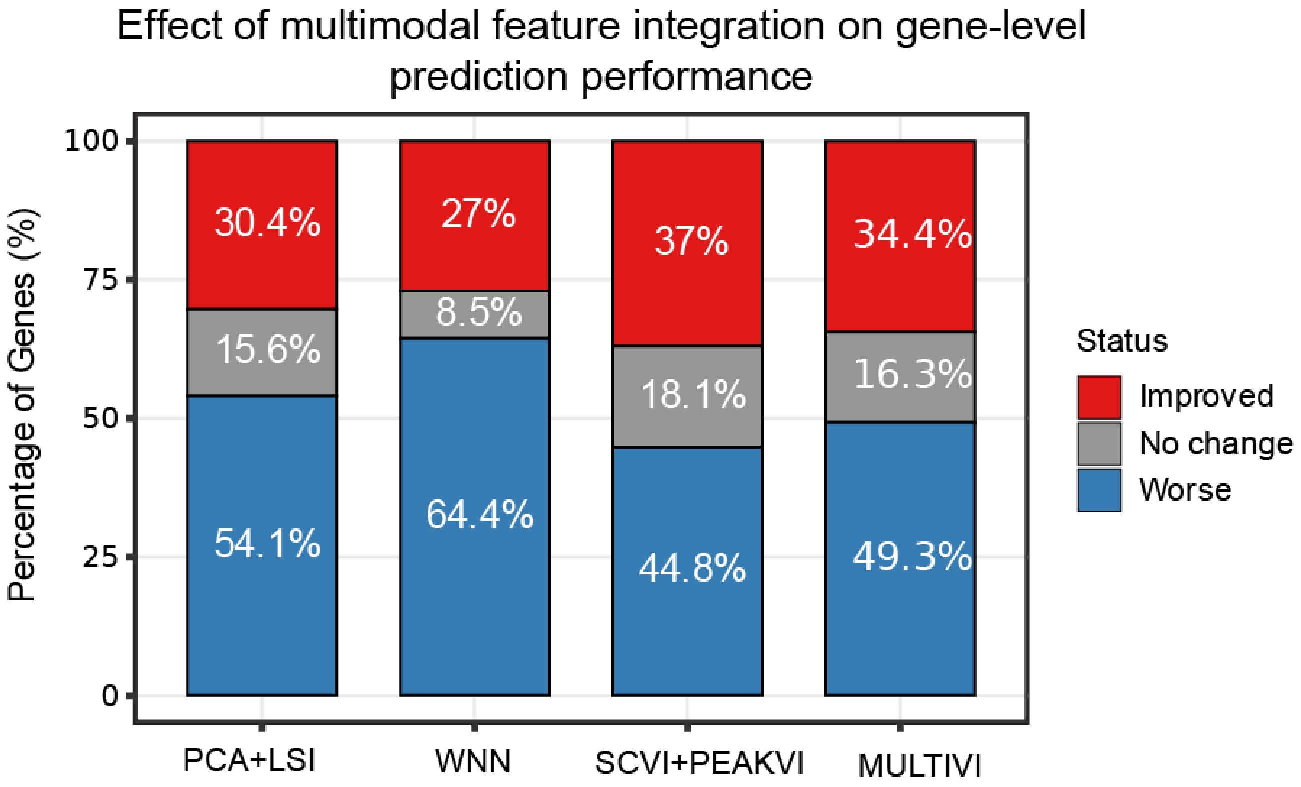


**Supplementary Figure S7: Gene-level impact of multimodal integration relative to RNA-only prediction.** Target genes were classified as improved, unchanged, or worsened based on the change in Spearman correlation (ΔSpearman) between multimodal and RNA-only models. Bars show the fraction of genes in each category for each integrated analysis strategy.

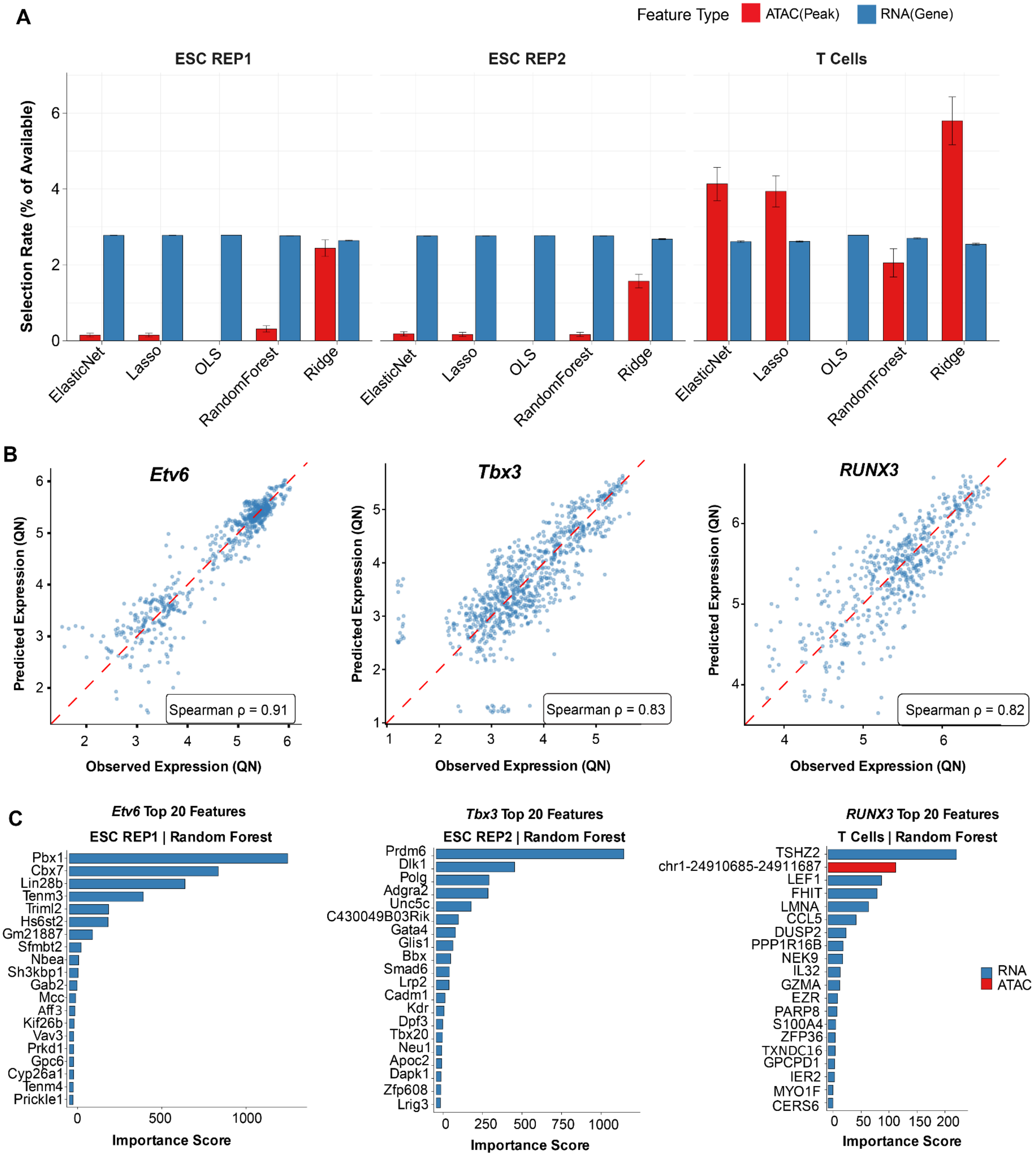


**Supplementary Figure S8: Feature interpretability of multimodal gene expression prediction.** (A) Selection rate of RNA-derived and ATAC-derived features across models and datasets. (B) Representative target gene predictions showing correlation between observed and predicted expression. (C) Top 20 predictive features identified by random forest importance scores for representative genes across datasets, highlighting contributions from RNA and ATAC features. QN:Quantile Normalized.

#### Supplementary Tables

**Table S2 | Gene-level classification of multimodal feature integration effects.** List of target genes classified as improved, unchanged, or worsened based on change in Spearman correlation between multimodal and RNA-only prediction. Classifications are reported for each dataset and integrated analysis strategy and correspond to the summary proportions shown in Supplementary Figure S7.
